# Synaptic Plasticity in the Agranular Insular Cortex Predicts Escalated Ethanol Consumption

**DOI:** 10.1101/2021.05.17.444509

**Authors:** Joel E. Shillinglaw, Heather C. Aziz, Daniela G. Carrizales, Richard A. Morrisett, Regina A. Mangieri

## Abstract

The Agranular Insular Cortex (AIC) is implicated in alcohol use disorder and pharmacologically relevant concentrations of acute ethanol inhibit N-methyl-D-aspartate receptor (NMDAR)-mediated glutamatergic synaptic transmission and plasticity onto layer 2/3 AIC pyramidal neurons. However, it is not known whether the actions of ethanol on glutamatergic synapses are means by which chronic ethanol alters mechanisms of learning and memory in AIC as alcohol drinking transitions from controlled to problematic. We utilized the chronic intermittent ethanol (CIE) vapor model of ethanol exposure in adult male mice, alone or in combination with voluntary ethanol consumption, to determine whether glutamatergic synapses on layer 2/3 AIC pyramidal neurons are differentially regulated by different durations and intensities of chronic ethanol exposure. We observed evidence of both ethanol- and age-related metaplasticity of AIC layer 2/3 glutamatergic synapses, as only young adult, ethanol-naïve mice exhibited NMDAR-dependent long term depression ex vivo. Our findings also indicated that voluntary ethanol consumption alone can elicit glutamatergic plasticity in vivo. We found that the ratio of NMDAR- to AMPAR-mediated postsynaptic currents was reduced not only in CIE-treated, but also in air-treated, chronically drinking mice relative to ethanol-naïve controls. Furthermore, lower NMDA/AMPA ratios were predictive of greater escalation of ethanol consumption. These findings suggest that even moderate exposure to ethanol may elicit plasticity in the agranular insular cortex that contributes to the progression toward uncontrolled drinking.

## Introduction

The Agranular Insular Cortex (AIC) is involved in interoceptive processing, and deficits in AIC functioning and interoceptive processing are implicated in alcohol use disorder (AUD)^1–3^. Despite evidence for altered insular functioning and/or output in animal models of AUD^4–8^, there has been minimal investigation of the effects of ethanol on synaptic transmission in the AIC^9,10^. It is axiomatic that synapses are ethanol-sensitive targets mediating the behavioral responses to ethanol, but the modulatory actions of ethanol on synaptic glutamate and gamma-aminobutyric acid (GABA) transmission have proven to vary by brain region. Thus, electrophysiological investigation of the effects of ethanol on the AIC is essential to uncover how AIC dysfunction occurs in the context of AUD.

We recently demonstrated that components of glutamatergic transmission are targets for the acute actions of ethanol in the AIC^9^. Specifically, we found that N-methyl-D-aspartate receptor (NMDAR)-mediated synaptic transmission and NMDAR-dependent plasticity onto AIC layer 2/3 pyramidal neurons were inhibited by acute ethanol at concentrations achieved during alcohol drinking. Prior work in several brain regions has shown that chronic ethanol produces long-term alterations in the synaptic targets of acute ethanol and that this ethanol-induced synaptic plasticity contributes to altered brain function as well as neural and behavioral components of AUD^11,12^. Therefore, we hypothesized that chronic ethanol exposure impairs AIC functioning by inducing glutamatergic synaptic adaptations.

To test this hypothesis, we utilized the chronic intermittent ethanol (CIE) vapor paradigm to model in vivo ethanol exposure in mice. CIE is a passive exposure model that yields stable intoxicating blood alcohol concentrations over extended periods of time (16 hours per day), and mimics a pattern of repeated cycles of binge intoxication and withdrawal^13,14^. Prior research from our laboratory observed that limited exposure to ethanol achieved via a single 4-day CIE treatment induces robust alterations in the polarity of NMDAR-dependent glutamatergic synaptic plasticity of medium spiny neurons in the nucleus accumbens shell in a cell-type specific manner^15,16^. Since repeated cycles of CIE result in enhanced volitional ethanol consumption by rodents^13,14,17–19^, these results are taken to indicate that even limited CIE exposure is sufficient to alter glutamatergic synaptic plasticity mechanisms, which may contribute to altered brain function and AUD-related pathology^20^. Here we used both limited and extended CIE treatment, alone (limited CIE experiment) or in conjunction with volitional consumption of ethanol (extended CIE experiments), to evaluate how chronic ethanol exposure affects glutamate synapses on layer 2/3 AIC pyramidal neurons. We found that repeated exposure to ethanol can impair NMDAR-mediated plasticity in the AIC, and that even volitional ethanol drinking is sufficient to induce glutamatergic synaptic plasticity in this region.

## Materials and Methods

### Mice

Adult (≥8 weeks old) *Drd1a*-tdTomato BAC transgenic male mice^21^, either hemizygous or null carriers of the transgene, were used for all experiments. A colony of *Drd1a*-tdTomato mice (initial breeding pairs obtained from The Jackson Laboratory, (IMSR Cat# JAX:016204, RRID:IMSR_JAX:016204) was maintained in the lab using matings in which only one parent carried the *Drd1a*-tdTomato transgene^22^. Mice were housed in standard cages (7.5” x 11.5” x 5”) with Sani-Chips wood bedding (PJ Murphy) at ∼22°C with a 12:12 light:dark cycle (lights on, Zeitgeber time 0 (ZT0), at 9:30 p.m.). Cages contained environmental enrichment in the form of a cotton fiber nestlet or a plastic hut. Water and standard chow (LabDiet®5LL2 Prolab RMH1800) were available ad libitum. All experimental procedures were approved by The University of Texas at Austin Institutional Animal Care and Use Committee, and were in accordance with the Guide for the Care and Use of Laboratory Animals as adopted by the National Institutes of Health.

### Chronic intermittent ethanol vapor exposure and two-bottle choice drinking

Chronic intermittent ethanol (CIE) vapor exposure, with and without two-bottle choice (2BC) ethanol drinking, was performed as previously described^15,17,19^. Briefly, 95% ethanol (Pharmco-AAPER) was placed inside a sealed flask and volatized by bubbling with air, supplied by an aquarium pump at a flow rate ∼200-300 mL/min. The resulting ethanol vapor was combined with an additional stream of air (∼3.5 L/min flow rate) and delivered to mice in vapor chamber units consisting of an airtight top, a vapor inlet, and an exhaust outlet (Allentown Inc., Allentown, NJ). Mice received intraperitoneal injections of ethanol (20% v/v) with pyrazole (1 mM) in phosphate buffered saline (PBS) immediately prior to placement in chamber units. At the start of each experiment, the loading dose was equivalent to 1.5 g/kg ethanol, 68.1 mg/kg pyrazole, but injection volumes (doses) and/or ethanol vapor flow rates were adjusted as needed to keep mice in the target BEC range of 150-200 mg/dL. CIE mice were exposed to ethanol vapor for 16 hours (from ZT19.5 to ZT11.5 of the following day), followed by 8 hours of withdrawal, during which mice were removed from the vapor chambers and not exposed to ethanol. Each “bout” of CIE consisted of four consecutive days of vapor exposure (16 hours on/8 hours off). Air-treated mice were handled identically but injected with a solution of pyrazole (1 mM, 68.1 mg/kg) in PBS before being placed into vapor chamber units that received only air (flow rate ≈ 3.5 L/min).

The limited CIE experiment (Figure 1) utilized one bout of CIE (or air) treatment and mice were euthanized 24 hours after removal from chambers to prepare brain slices for electrophysiology. For extended CIE experiments (Figures 2, 4, 5, and 6), mice received three bouts of CIE (or air) treatment in conjunction with 2BC. The first bout of chambering was preceded by 21 days of daily, 2-hour, 2BC drinking sessions in which home cage water bottles were replaced (from ZT11.5 to ZT13.5) with two bottles containing either tap water or 15% ethanol (Pharmco-AAPER) in tap water (vol/vol). Bottles were weighed immediately before and after each session and the difference in weight was used to calculate consumption. The first and second bouts of chambering were followed by 3 days in which mice were left undisturbed and then 5 days of 2BC sessions. Following the third bout of chambering, mice were left undisturbed for 5-8 days and then were euthanized to prepare brain slices for electrophysiology. This duration of withdrawal from ethanol was selected because it corresponds to a period in which mice reliably exhibit CIE-induced enhancement of volitional ethanol consumption in our hands^17,19^.

**Figure 1.**
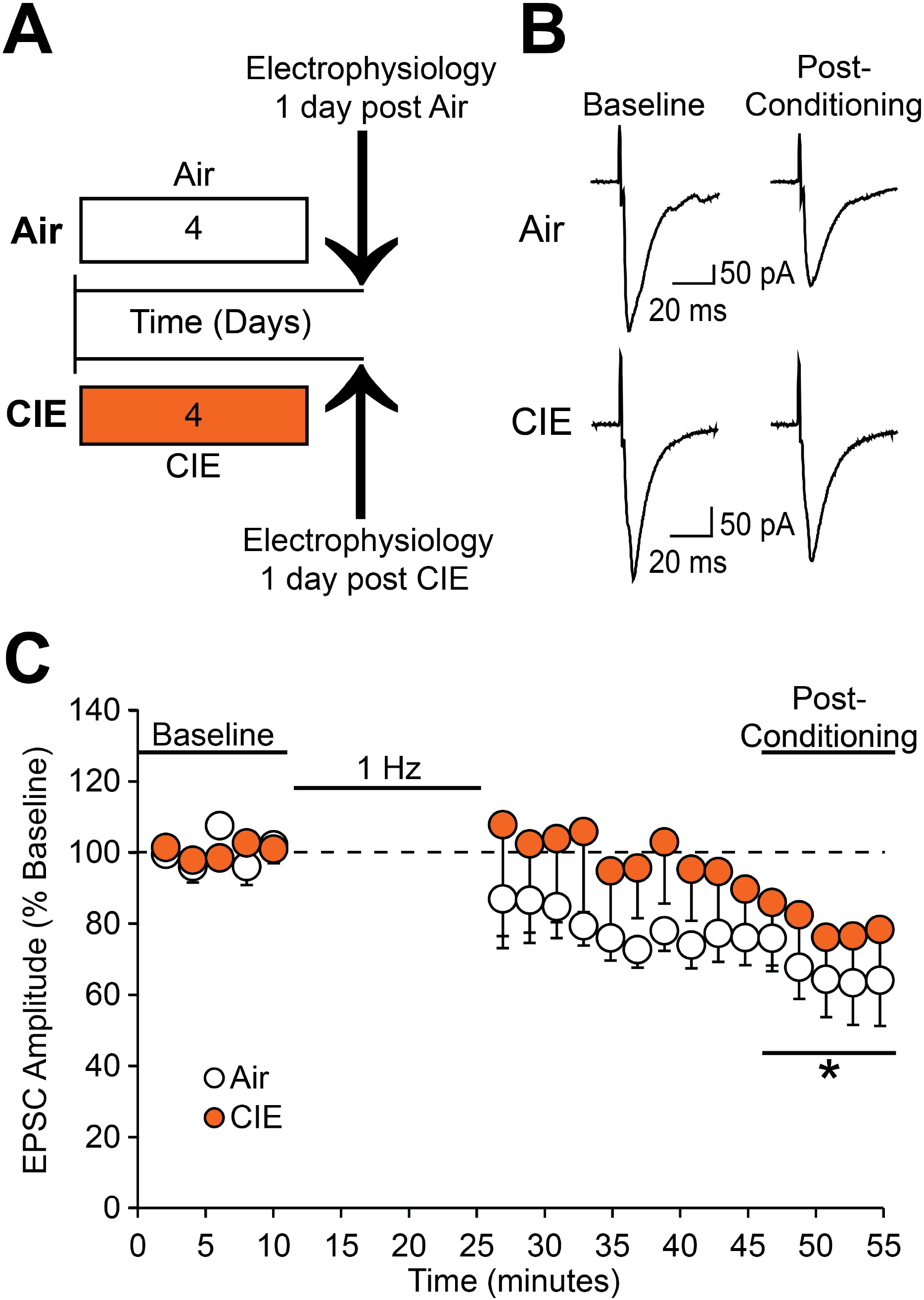
Limited exposure to CIE disrupts LTD expression at glutamatergic synapses onto AIC layer 2/3 pyramidal neurons. (A) Timeline for limited CIE (or air) exposure prior to ex vivo electrophysiological recordings. (B) Representative traces from a single neuron of each group showing evoked EPSCs before (“Baseline”), and 20-30 min after (“Post-Conditioning”), stimulation protocol (900 pulses at 1 Hz while holding the neuron at −70 mV). (C) Conditioning stimulation induced LTD in Air-treated (n = 8 neurons/8 slices/8 mice; *, p = 0.014 post-conditioning period versus baseline), but not CIE-treated mice 24 hours into withdrawal (n = 10 neurons/8 slices/8 mice). Values are expressed as averages ± S.E.M.

**Figure 2.**
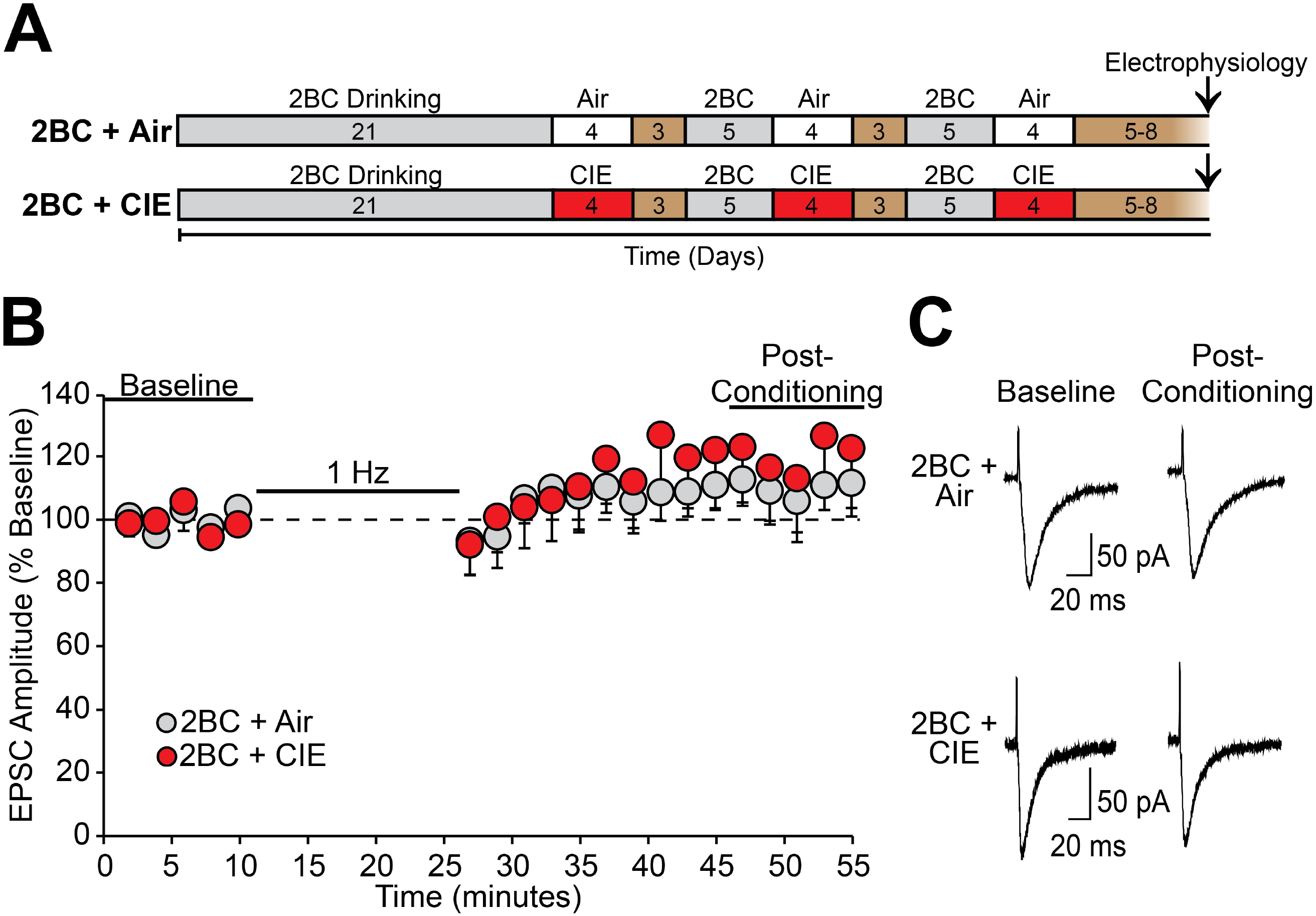
Extended CIE exposure with ethanol drinking, and extended air exposure with ethanol drinking, are accompanied by an absence of LTD expression in AIC layer 2/3 pyramidal neurons. (A) Timeline for extended CIE (or air) exposure and two-bottle choice (2BC) ethanol drinking (15% ethanol or water) prior to ex vivo electrophysiological recordings. Brown fill indicates days when mice were left unperturbed in home cages. (B) Conditioning stimulation did not induce LTD of evoked EPSCs onto AIC layer 2/3 pyramidal neurons of 2BC + Air (n = 8 neurons/8 slices/6 mice) or 2BC + CIE (n = 8 neurons/8 slices/7 mice) mice. (C) Representative traces from a single neuron of each group showing evoked EPSCs before (“Baseline”), and 20-30 min after (“Post-Conditioning”), stimulation protocol. Values are expressed as averages ± S.E.M.

**Figure 3.**
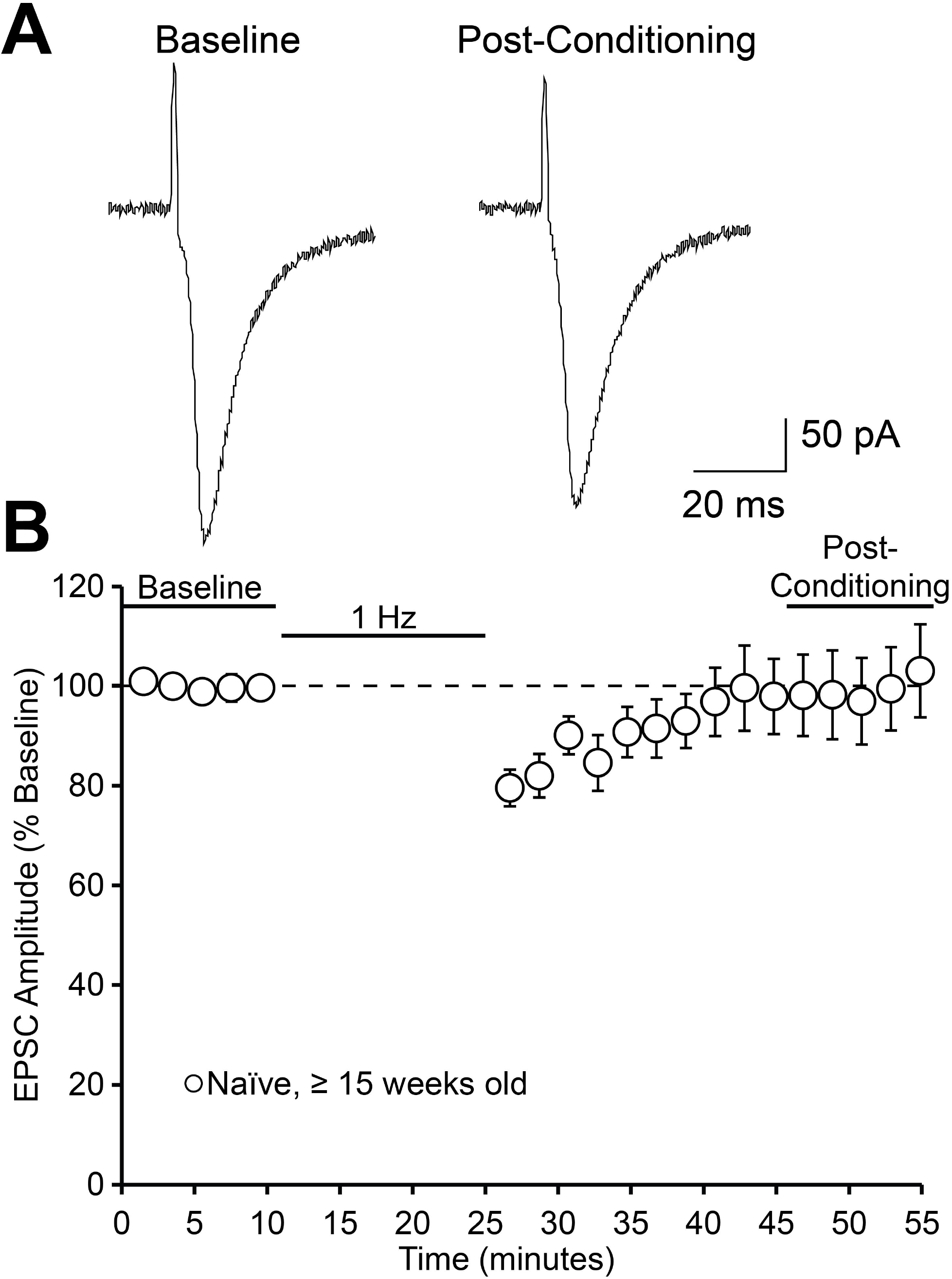
LTD at glutamatergic synapses onto AIC layer 2/3 pyramidal neurons is not observed in slices from ethanol-naïve mice that are at least 15 weeks old. (A) Representative traces from a single neuron showing evoked EPSCs before (“Baseline”), and 20-30 min after (“Post-Conditioning”), stimulation protocol. (B) Conditioning stimulation did not induce LTD of evoked EPSCs onto AIC layer 2/3 pyramidal neurons from mice that were at least 15 weeks old (n = 12 neurons/12 slices/8 mice). Values are expressed as averages ± S.E.M.

**Figure 4.**
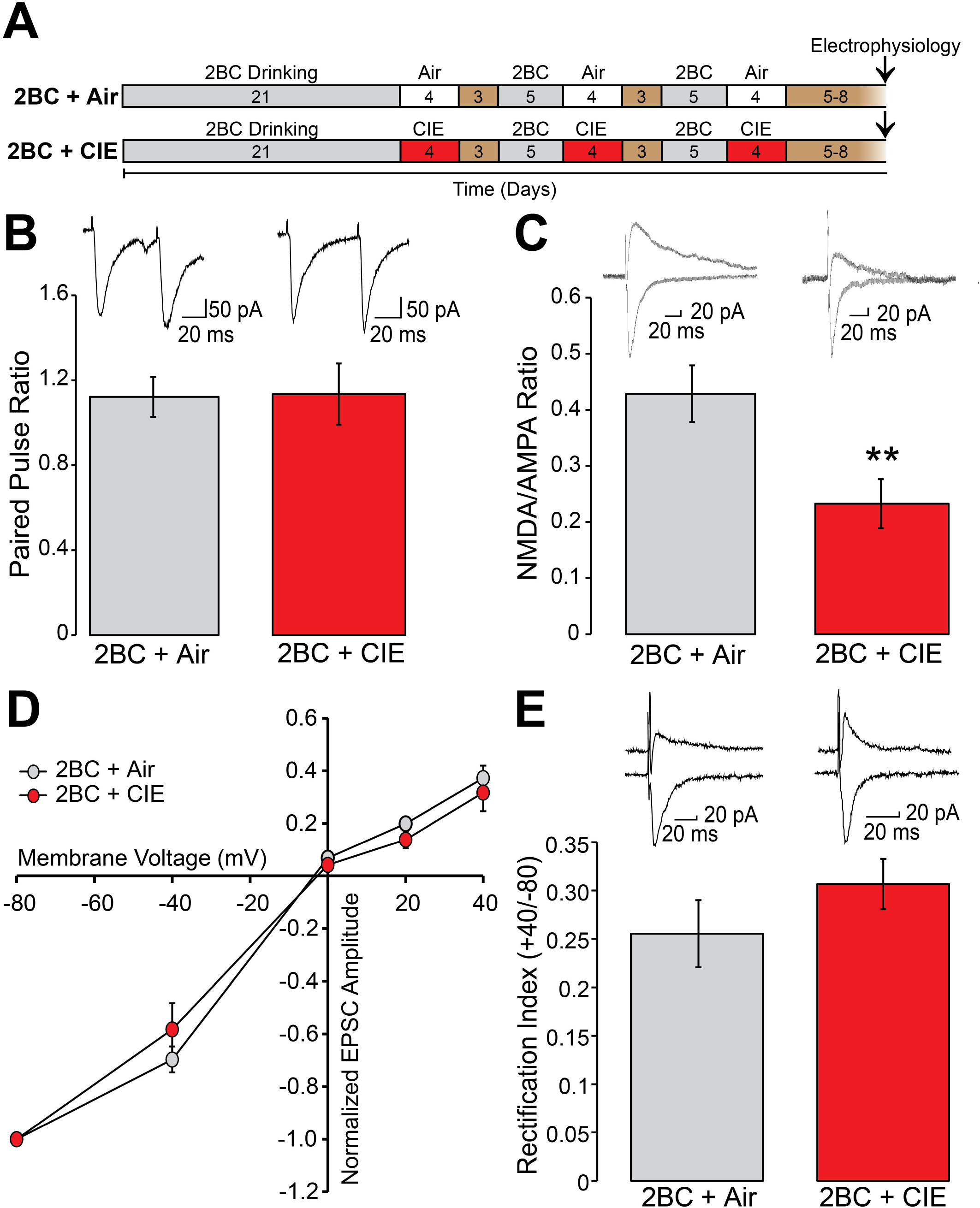
Extended exposure to CIE and ethanol drinking reduces NMDA/AMPA ratio in AIC layer 2/3 pyramidal neurons, but does not alter the expression of other glutamatergic electrophysiological properties, relative to an air exposed, ethanol drinking group. (A) Timeline for extended CIE (or air) exposure and two-bottle choice (2BC) ethanol drinking (15% ethanol or water) prior to ex vivo electrophysiological recordings. Brown fill indicates days when mice were left unperturbed in home cages. (B) Paired pulse ratio (second evoked EPSC amplitude/ first evoked EPSC amplitude) did not differ between 2BC + Air (n = 8 neurons/7 slices/7 mice) and 2BC + CIE mice (n = 9 neurons/8 slices/4 mice). (C) NMDA/AMPA ratio was decreased in 2BC + CIE (n = 9 neurons/9 slices/5 mice) relative to 2BC + Air mice (n = 9 neurons/9 slices/5 mice). **, p = 0.0098, 2BC + CIE vs. 2BC + Air. (D) Current-voltage relationship of AMPAR-mediated EPSCs for 2BC + Air (n = 10 neurons/10 slices/6 mice) and 2BC + CIE mice (n = 6 neurons/6 slices/4 mice). (E) No difference in the rectification index (ratio of EPSC amplitudes evoked at +40 mV and −80 mV holding potentials) was observed between 2BC + Air (n = 13 neurons/13 slices/6 mice) and 2BC + CIE mice (n = 16 neurons/15 slices/8 mice). Values are expressed as averages ± S.E.M.

**Figure 5.**
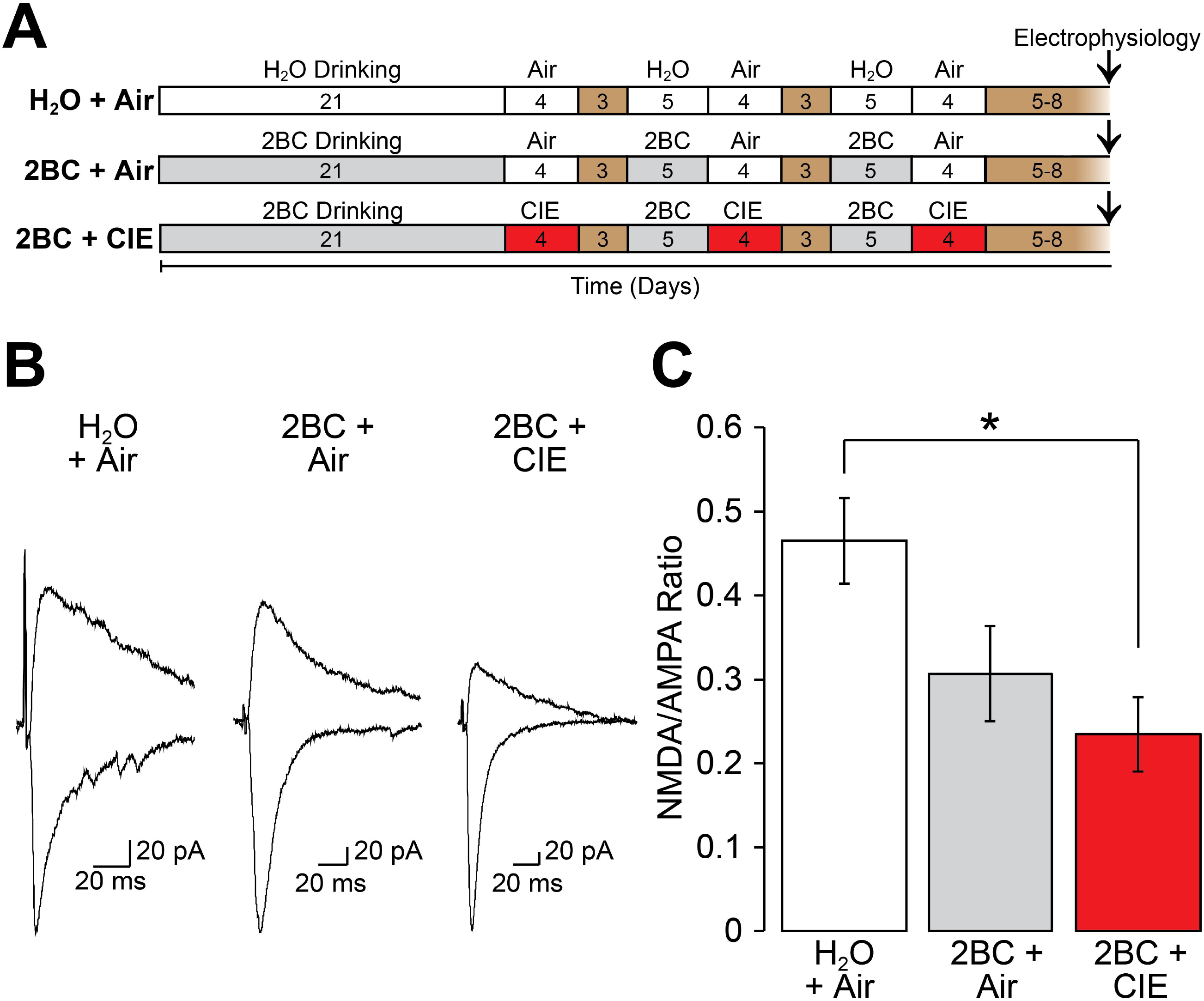
Extended ethanol exposure reduces NMDA/AMPA ratio in AIC layer 2/3 pyramidal neurons. (A) Timeline for extended CIE (or air) exposure with two-bottle choice ethanol drinking (15% ethanol or water), or extended air exposure with water drinking (handling control group, “H_2_O-Air”), prior to ex vivo electrophysiological recordings. Brown fill indicates days when mice were left unperturbed in home cages. (B) Representative traces from a single neuron of each group showing evoked NMDA/AMPA ratio. (C) NMDA/AMPA ratio was altered by ethanol exposure (p = 0.011, 1-way ANOVA) and significantly decreased in 2BC + CIE mice relative to H_2_O + Air mice (*, p = 0.014, Bonferroni post-hoc comparison). H_2_O + Air, n = 14 neurons/9 slices/5 mice; 2BC + Air, n = 8 neurons/5 slices/2 mice; 2BC + CIE, n =8 neurons/4 slices/3 mice.

**Figure 6.**
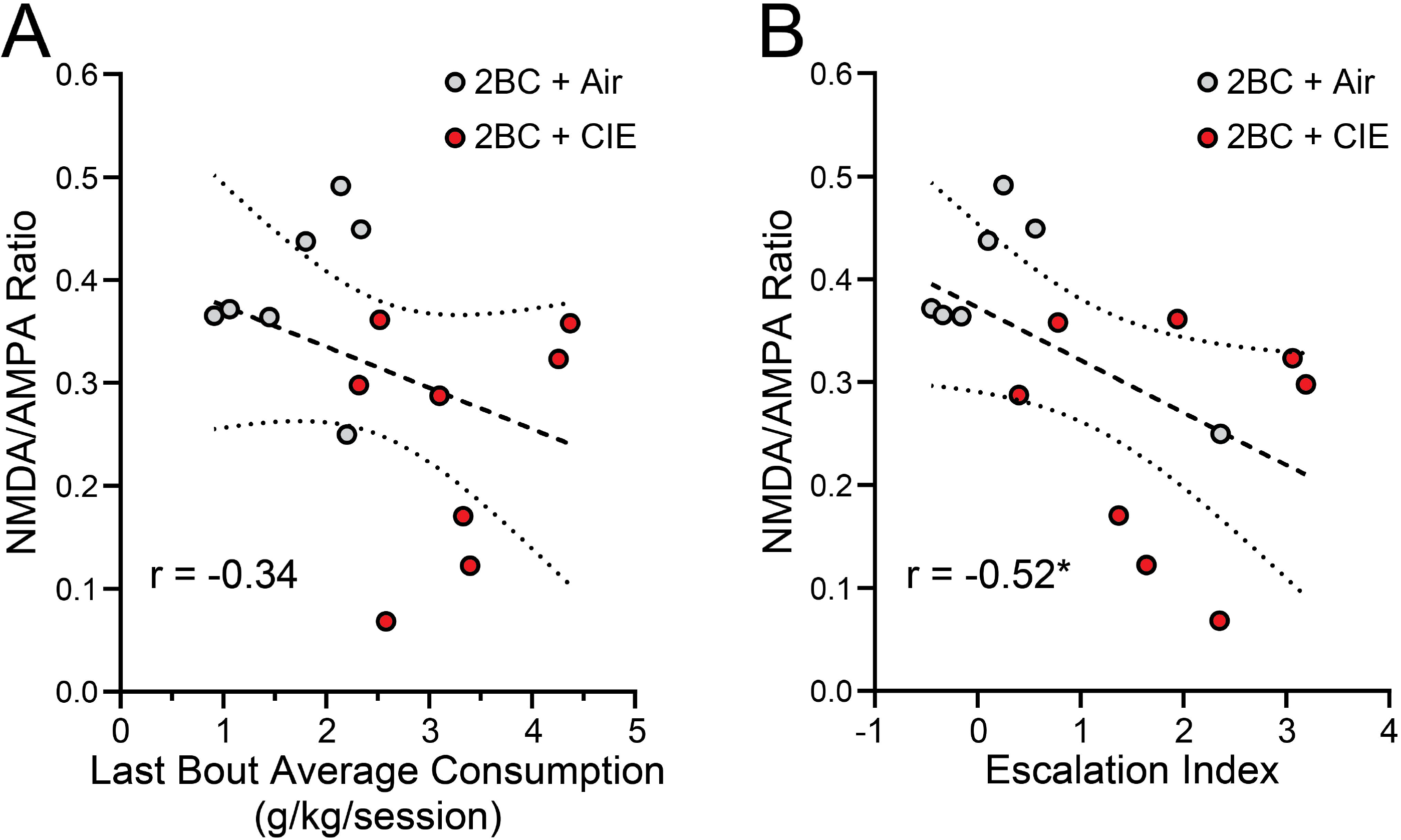
Relationships between ethanol drinking measures and NMDA/AMPA ratio in AIC layer 2/3 pyramidal neurons. Data from the 2BC + CIE and 2BC + Air groups shown in Figures 4 and 5 were combined and an average NMDA/AMPA ratio was calculated for each mouse. (A) NMDA/AMPA ratio versus average ethanol consumption (g ethanol per kg body weight) during the final five 2BC sessions. (B) NMDA/AMPA ratio versus escalation index. The escalation index expresses each animal’s change in ethanol consumption relative to its initial consumption, and was calculated as (final g/kg – initial g/kg)/initial g/kg using 5-session averages of the final five and initial five 2BC sessions. Dashed and dotted lines indicate plots of the best-fit lines and 95% confidence intervals, respectively. Correlation coefficients are indicated on each graph. *, p = 0.02, Pearson r.

In one of the extended CIE experiments (Figure 5), a handling control group was concurrently used. These mice were handled identically to mice in the other groups, but they remained ethanol naïve; that is, they underwent 2BC drinking sessions with water in both bottles and received air-treatment procedures during chambering.

### Blood ethanol concentrations

Immediately upon removal from vapor chambers each day, tail blood samples were taken and blood ethanol concentrations (BECs) were determined using a gas chromatograph (Bruker 430-GC) equipped with a flame ionization detector and CombiPAL autosampler (Bruker Corporation, Fremont, CA). Mice were briefly restrained and two, 5 µL, tail blood samples were collected. Each sample was immediately pipetted into a 10 mL vial containing 45 µLs of saturated sodium chloride solution. Vials were heated to 65 °C, and ethanol vapor was absorbed by a solid-phase micro extraction fiber (SPME; 75 µm CAR/PDMS, fused silica; Supelco, Bellefonte, PA). A capillary column was the stationary phase (30 m x 0.53 mm x 1 µm film thickness; Agilent Technologies, Santa Clara, CA), and helium was used as the mobile phase (at a flow rate of 8.5 mL/min). CompassCDS Workstation software (Bruker Corporation, Fremont, CA) was used to analyze ethanol peaks, and external ethanol standards were included in every run to construct standard curves for interpolation of ethanol concentrations.

### Preparation of Brain Slices

Mouse brains were rapidly extracted and coronal slices containing the most anterior portion of the AIC were collected via established laboratory protocols^9^. Briefly, mice were first anesthetized by inhalation of isoflurane and then euthanized by decapitation. Brains were then quickly extracted from the skull and submerged in ice-cold oxygenated artificial cerebrospinal fluid (ACSF) containing the following (in mM): 210 Sucrose, 26.2 NaHCO3, 1 NaH2PO4, 2.5 KCl, 11 dextrose, bubbled with 95% O2/5% CO2. Coronal slices (250 to 270 µm thick) containing AIC were collected using a Leica VT1000S vibrating microtome (Leica Corp., Bannockburn, IL), which were then placed into an incubation solution containing the following (in mM): 120 NaCl, 25 NaHCO3, 1.23 NaH2PO4, 3.3 KCl, 2.4 MgCl2, 1.8 CaCl2, 10 dextrose, continuously bubbled with 95% O2/ 5% CO2; 32°C, for at least 45 min for recovery prior to recording.

### Patch-Clamp Electrophysiology

As previously described^9^, whole cell recordings were taken from layer 2/3 pyramidal AIC neurons identified via morphology (large, pyramidal-like shape) using a MRK200 Modular Imaging system (Siskiyou Corporation, Grants Pass, OR) mounted on a vibration isolation table. Recordings were made in ACSF containing (in mM): 120 NaCl, 25 NaHCO3, 1.23 93 NaH2PO4, 3.3 KCl, 1.2 MgSO4, 2.0 CaCl2, and 10 dextrose unless otherwise noted, bubbled with 95% O2/ 5% CO2; 32°C, controlled by an in-line heater (Warner Instruments, Hamden, CT). Picrotoxin (50 µM) was included in the recording ACSF to block GABA_A_ receptor-mediated synaptic currents. Brain slices were perfused at a rate of 2.0 mL/min. A P-97 Flaming/Brown model micropipette puller (Sutter Instruments, San Rafael, CA) was used to make recording electrodes (thin-wall glass, WPI Instruments, Sarasota FL) of resistances from 3-6 MΩ. For all experiments, recording electrodes were filled with (in mM): 120 CsMeSO4, 15 CsCl, 8 NaCl, 10 HEPES, 0.2 EGTA, 10 TEA-Cl, 4 Mg-ATP, 0.3 Na-GTP, 0.1 spermine, and 5 QX-314-Cl. Series resistance (Rs) and holding current (I_hold_) were monitored throughout each experiment. Cells with Rs of over 30 MΩ or that changed over 20% over the course of the experiment were excluded from the analysis. I_hold_ was also used to identify recordings with significant instability of the patch and recordings with I_hold_ in excess of −500 pA for several minutes were excluded. One cell per brain slice was used for LTD experiments, and multiple cells per brain slice were sometimes used for other glutamatergic assays. All electrophysiology chemicals were obtained from either Fisher Scientific, Sigma-Aldrich, or Tocris Bioscience.

An Axopatch 200B amplifier (Axon Instruments, Foster City, CA) was used to acquire all currents, which were filtered at 1 kHz, and digitized at 10-20 kHz via a Digidata 1440A interface board using pClamp 10.2 (Axon Instruments). Excitatory postsynaptic currents (EPSCs) were evoked with a stainless steel bipolar stimulating electrode (MX21AES, FHC, Inc., Bowdoin, ME, United States) placed approximately 500 µm dorsomedial to the cell body. For LTD experiments, EPSCs were evoked for at least 10 min (at 0.025 Hz) to ensure stable recordings, followed by a low-frequency stimulation protocol consisting of 1 Hz stimulation for 15 minutes. Evoked EPSCs were then monitored for a 30 minute post-stimulation period at 0.025 Hz to test for the expression of LTD. Neurons were held at −70 mV for the entirety of LTD experiments. EPSC amplitudes were normalized by dividing each amplitude by the average baseline amplitude and the % baseline values were used for statistical analysis. The time courses for LTD experiments are presented with normalized data averaged in 2 min bins. Paired-pulse ratios were determined by applying two stimuli of equal intensity, separated by an interstimulus interval of 50 ms and calculated as the ratio of EPSC amplitudes (EPSC 2/EPSC 1). NMDA/AMPA ratios were calculated as the ratio of the EPSC evoked at +40 mV holding potential, amplitude measured 50 ms after stimulation (NMDA), and the peak amplitude of the EPSC evoked at −80 mV (AMPA). Current-voltage (I-V) relationships and rectification of AMPAR-mediated currents were determined by stepping the command voltage over a range of potentials and evoking EPSCs in the presence of 100 µM DL-APV. Rectification index was calculated as the EPSC amplitude evoked at +40 mV/EPSC amplitude at −80 mV.

### Data Analysis

GraphPad Prism version 8.0 or higher was used to perform statistical analyses. Expression of LTD was determined by comparing the average normalized evoked EPSC amplitude during the 20 to 30 minute period after the low-frequency stimulation protocol to the 10 minute baseline period using a one-sample t test with the theoretical mean equal to 100. Group comparisons of paired pulse ratio, NMDA/AMPA ratio, and rectification index were made using unpaired t-tests or 1-way ANOVA with Bonferroni post hoc tests, as appropriate. AMPAR I-V curves were compared between groups using an F-test to compare the linear fits of each curve. Relationships between ethanol consumption measures and NMDA/AMPA ratio were assessed using simple linear regression and Pearson correlation analyses. Statistical significance was defined as p < 0.05.

## Results

### Expression of LTD following limited or extended ethanol experience

We first tested whether limited CIE (1-bout) affected the expression of AIC glutamatergic LTD. Brain slices containing the AIC were prepared from mice 24 hours after limited (1-bout) CIE or air treatment, and layer 2/3 AIC neurons were tested for their response to a low frequency stimulation paradigm (1 Hz for 15 min) previously shown to induce NMDAR-dependent LTD of evoked EPSCs^9^. We found that LTD was produced in neurons from ethanol-naïve air-treated mice (Figure 1; one-sample t-test, t(7) = 3.3, p = 0.014). However, using this same stimulation paradigm, we were unable to induce LTD in neurons from CIE mice 24 hours after the last ethanol vapor exposure (Figure 1; one-sample t-test, t(9) = 1.5, p = 0.17).

In the next set of experiments, we used an extended CIE protocol, which has been shown to enhance volitional ethanol drinking and produce ethanol dependence and withdrawal^13,14,18^. Mice were exposed to three bouts of either CIE or air treatment, with each bout preceded by 2BC drinking sessions (2BC + CIE and 2BC + Air groups, respectively; Figure 2A). Glutamatergic synaptic events were recorded from layer 2/3 AIC neurons ex vivo 5-8 days after the final CIE or air bout, a time period that corresponds to the point at which CIE-treated mice consistently exhibit enhanced 2BC ethanol drinking relative to air-treated mice^17,19^. As we observed following limited CIE exposure, neurons from 2BC + CIE mice did not exhibit LTD of evoked EPSCs in response to 1 Hz conditioning stimulation (one-sample t-test, t = 1.1, p = 0.29; Figure 2B,C). We also did not observe LTD in neurons from 2BC + Air mice (one-sample t-test, t = 1.2, p = 0.28; Figure 2B,C).

At face value, this pattern of results might seem to suggest that even volitional exposure to ethanol via 2BC drinking sessions is sufficient to disrupt LTD in layer 2/3 AIC neurons. However, the failure to observe LTD in the 2BC + Air group was unexpected, because similar studies of NMDAR-LTD have found that this form of LTD was still observable following 2BC + Air treatment^17,19^. NMDAR-LTD in the AIC has not been extensively characterized, but, in general, synaptic physiology and expression of plasticity can change from adolescence to adulthood, and even beyond the emergence of early adulthood^23–27^. Our prior work demonstrating the existence of this form of plasticity in the AIC^9^, and the limited CIE experiment of the present work, used animals that were younger in age (ranging from ∼7-13 weeks) than those of the extended ethanol exposure experiment (≥ 15 weeks). Thus, an alternative explanation for the absence of LTD in the 2BC + Air group of the extended ethanol exposure experiment might be the older age of the mice. To examine this possibility, we determined whether we could induce LTD of evoked EPSCs in AIC layer 2/3 neurons from ethanol-naïve mice that were at least 15 weeks of age, using the same low frequency stimulation paradigm as before. Under these conditions, we did not observe LTD (Figure 3; one-sample t-test, t(11) = 0.06, p = 0.96). This finding suggests that the expression of this form of plasticity is an age-dependent phenomenon in layer 2/3 AIC neurons, and, furthermore, indicates that this assay cannot be used to evaluate ethanol-elicited synaptic adaptations in mice that are 15 weeks or older.

### Glutamatergic pre- and post-synaptic properties following extended ethanol experience

Prior studies in other brain regions showing changes in the ability to induce LTD have suggested that this observation may indicate other phenomena, such as a change in NMDAR function or an increase in the proportion of calcium-permeable, to calcium-impermeable, AMPA receptors^19,28^. Therefore, we also determined whether 2BC + CIE differentially modulated other indices of glutamatergic synaptic transmission and plasticity relative to 2BC + Air (Figure 4A). We found no difference in the paired pulse ratio between 2BC + CIE and 2BC + Air mice (Figure 4B; unpaired t-test, t(15) = 0.07, p = 0.94). We observed a significant reduction in the NMDA/AMPA ratio in the 2BC + CIE relative to the 2BC + Air mice (Figure 4C; unpaired t-test, t(16) = 2.9, p = 0.0098). Finally, the current-voltage relationship (Figure 4D; F test, F(2,76) = 0.72, p = 0.49) and the rectification index for evoked AMPAR currents (Figure 4E; unpaired t-test, t(27) = 1.21, p = 0.24) were not different between 2BC + CIE and 2BC + Air mice.

### Ethanol drinking alone suppresses NMDA/AMPA

The reduced NMDA/AMPA ratio in the 2BC + CIE group, relative to the 2BC + Air group, suggests that this alteration in AIC function may relate to the progression toward pathological ethanol drinking. We wondered, though, whether drinking alone (without CIE) can impact NMDA/AMPA. Therefore, we replicated the extended CIE experiment with additional groups of 2BC + Air and 2BC + CIE mice, along with a concurrent ethanol-naïve handling control group, “H_2_O + Air”, which was treated identically to the 2BC + Air group, but received water in both bottles during 2BC sessions (Figure 5A). In this set of experiments, we observed a main effect of group on the NMDA/AMPA ratio in layer 2/3 AIC neurons (one-way ANOVA, F(2, 27) = 5.4, p = 0.011), but the difference in NMDA/AMPA ratios between 2BC + Air and 2BC + CIE groups was not statistically significant (p = 0.13, Bonferroni’s multiple comparisons test); Figure 5B,C. The results of both extended exposure experiments taken together strongly indicate that chronic ethanol experience suppresses the NMDA/AMPA ratio in layer 2/3 AIC pyramidal neurons, but this may not require CIE-induced dependence and can even occur in response to more moderate exposure to ethanol. To further examine the relationship between this glutamatergic adaptation and drinking behavior, we pooled the data from both experiments and assessed whether the NMDA/AMPA ratio was correlated with ethanol intake. We found that the relationship between an animal’s ultimate level of consumption and NMDA/AMPA ratio (Figure 6A) was modest and not statistically significant (Pearson’s r = −0.34, p = 0.10). However, the relationship between the degree to which an animal escalated in consumption of ethanol over the course of the experiment and the NMDA/AMPA ratio was evident (Pearson’s r = −0.52, p = 0.02), with escalation of drinking being inversely correlated to NMDA/AMPA ratio; Figure 6B.

## Discussion

Chronic ethanol exposure disrupts the normal functioning of prefrontal cortical regions via long-term adaptations of neuronal electrophysiological activity^29–34^. These neuroadaptations may reveal, at least in part, how various cortically mediated cognitive-behavioral processes, such response inhibition and reversal learning, are disrupted in individuals with AUD^30,33^. Likewise, it has been shown that insula-dependent interoceptive processes are involved in aspects of AUD^1–3^, indicating that homeostatic physiology in this cortical region is also disrupted by chronic ethanol exposure. Our previous investigation of AIC layer 2/3 pyramidal neurons found that NMDAR-dependent transmission and glutamatergic synaptic plasticity are acutely disrupted by ethanol^9^. Therefore, the goal of the present work was to test whether chronic ethanol also alters AIC function via effects on glutamatergic synaptic transmission.

We found that following passive exposure to ethanol (limited CIE treatment) we did not observe LTD ex vivo when tested 24 hours into ethanol withdrawal under conditions that elicited NMDAR-dependent LTD in AIC neurons from control mice. Since the CIE vapor model of ethanol administration reproduces several behavioral and physiological aspects of AUD^18^, it is possible that disruption of NMDAR-dependent LTD may be a signature of altered AIC functioning as ethanol consumption shifts from controlled to problematic (AUD-like). Such a finding wound not be without precedence, as many groups have found, in other addiction-relevant brain regions, alterations in NMDAR-dependent plasticity that were linked to ethanol-related behaviors^17,22,35,36^. However, it is also possible that the observed disruption in AIC LTD has no relevance to ethanol-related behavior, and, rather, is a general consequence of ethanol withdrawal unrelated to the pathology of AUD. Indeed, Spiga and colleagues have suggested that the disruption of NMDAR-dependent LTD might be ubiquitous at excitatory synapses across several brain regions as a result of ethanol withdrawal^37^.

Therefore, we next employed additional bouts of CIE along with 2BC drinking to determine whether the disruption of AIC NMDAR-dependent LTD induced by passive exposure to high doses of ethanol is linked to ethanol-reinforced behavior. After extended ethanol experience, neither the 2BC + CIE nor the 2BC + Air group displayed LTD in response to conditioning stimulation. One possible interpretation of this finding is that NMDAR-LTD in the AIC is more sensitive to ethanol exposure than it is in other brain regions^17,19^, and that extended 2BC drinking alone was sufficient to disrupt LTD in this region. However, we sought to rule out other factors that could explain this unexpected result. Mice in the limited CIE experiment ranged from 8-11 weeks of age, while mice in extended exposure experiment were ≥15 weeks old. It is generally thought that adulthood starts for mice around postnatal day 60^38^, and so at first impression differences in mice from these different periods of adulthood may not appear important for experimental design. However, there is literature indicating that synaptic properties are at least partly age-dependent^23,24^ and can change even over the course of adulthood^25–27^. In our previous report, we were the initial group to demonstrate AIC NMDAR-dependent LTD using whole-cell voltage clamp electrophysiology in mice aged 7-13 weeks^9^ and we did not test whether this form of plasticity is age-dependent. Here we discovered that the same synaptic conditioning stimulation that elicits LTD in AIC layer 2/3 neurons from young adult mice (∼8-11 weeks old) did not do so in neurons from mice that were at least 15 weeks old. This suggests that the expression of NMDAR-dependent LTD in the AIC may be an age-dependent phenomenon, or that different stimulation parameters are required to induce LTD in older mice. In either case, the specific LTD assay we employed (which used 15 minutes of 1 Hz stimulation) appears to only be suitable for probing AIC glutamatergic adaptations in mice of a specific age range.

Since chronic ethanol has been shown to alter multiple aspects of glutamatergic synaptic transmission^12,39^, we evaluated whether extended CIE treatment of ethanol drinking mice affected measures of presynaptic glutamate release and post-synaptic signaling by AMPA and NMDA receptors, relative to air treatment of ethanol drinking mice. It is generally accepted that acute ethanol is an inhibitor of excitatory signaling, but chronic ethanol increases excitatory drive by increasing glutamatergic transmission onto neurons^11^. However, when comparing groups of drinking mice (2BC + CIE and 2BC + Air), we found no evidence that CIE affected presynaptic glutamate release probability onto AIC 2/3 pyramidal neurons as there was no difference in the paired pulse ratio. To assess postsynaptic function, we tested for a difference in the expression of calcium-permeable AMPA receptors (CPARS), which have been shown to be increased in several brain regions after exposure to ethanol and other drugs of abuse and are implicated in the escalation of seeking and intake of ethanol and other drugs^39^. We did not observe that CIE altered postsynaptic CPAR expression relative to air treatment, as AMPAR rectification index and I/V curves did not differ between groups.

We did, however, find in this set of experiments that the NMDA/AMPA ratio, a commonly used index of glutamatergic synaptic transmission and plasticity, was significantly reduced in 2BC + CIE mice relative to 2BC + Air mice. This observation prompted a subsequent set of experiments that included an ethanol-naïve handling control group, leading to our main finding – that drinking ethanol alone can alter the NMDA/AMPA ratio in layer 2/3 AIC pyramidal neurons. Since exposure to chronic ethanol has frequently been shown to lead to enhanced glutamatergic transmission via increased NMDAR function and NMDAR-mediated signaling^11,12,40,41^, we were initially surprised that ethanol exposure resulted in a suppressed NMDA/AMPA ratio in our experiments. Furthermore, although there is evidence that chronic ethanol also upregulates the expression and functioning of AMPA receptors, this generally has been shown to be at a lower level than that of NMDA receptors^11,12,42,43^. Therefore, at initial glance, our findings were perplexing as we expected a priori to observe an enhanced NMDA/AMPA ratio due to increased NMDAR expression and function accompanied by possible, but less significant, changes in AMPAR expression and function. However, it is becomingly increasingly clear that the effects of chronic ethanol on postsynaptic glutamate receptors depend upon the brain region investigated and the time point after ethanol withdrawal^30,33,44,45^. To better understand whether the difference in ratio we observed in our experiments resulted from changes in AMPARs, NMDARs, or both, additional follow-up electrophysiological assessments of isolated NMDAR-mediated and AMPAR-mediated input-output curves at this time point are warranted, as are biochemical approaches investigating whether changes in NMDAR and AMPAR subunit composition co-occur with this synaptic phenotype.

Finally, we found that although CIE vapor treatment tended to cause a greater suppression of the NMDA/AMPA ratio, ethanol drinking alone was sufficient to elicit this adaptation. The fact that the NMDA/AMPA ratio was suppressed in the 2BC + Air group, relative to ethanol naïve controls, indicates that glutamatergic synaptic changes in layer 2/3 AIC pyramidal neurons occur prior to the development of excessive drinking behavior. Interestingly, escalation of ethanol consumption, regardless of whether mice were treated with CIE, was correlated with plasticity as indexed by the NMDA/AMPA ratio – mice with lower ratios were those that had exhibited greater increases in the dose of ethanol self-administered over the course of the experiment. This set of findings, summarized in Figure 7, raises the question of whether even moderate ethanol exposure may have consequences that are relevant to other behavioral manifestations of cortical glutamatergic synaptic plasticity. For example, glutamatergic synaptic plasticity mechanisms in the insula have been shown to encode for phenotypes produced by models of neuropathic pain, as well as regulate the rate of conditioned taste aversion learning^46–48^. From this perspective, it would be an invaluable investigation to determine whether the glutamatergic synaptic changes in the 2BC + Air exposure paradigm are enough to modulate the expression of neuropathic pain as well as the extinction of conditioned taste aversion^46–48^. Moreover, ethanol has well-documented aversive properties, and the reduction in sensitivity to the aversive aspects of ethanol consumption have been increasingly recognized as a component of the addiction cycle for ethanol^49^. It has been shown that a single 24 hour experience in a two-bottle choice paradigm enhances consumption of ethanol adulterated with the bitter tastant quinine, but does not change quinine palatability^50^. Therefore, it could be that, in some mice, even early ethanol exposure or moderate doses of ethanol elicit glutamatergic synaptic plasticity in the insula and thereby disrupt interoceptive processing of aversive stimuli. Thus, one intriguing hypothesis suggested by our findings is that ethanol-elicited adaptation in AIC glutamatergic transmission may cause a suppression of aversive processing and thereby facilitate escalation of ethanol consumption.

**Figure 7.**
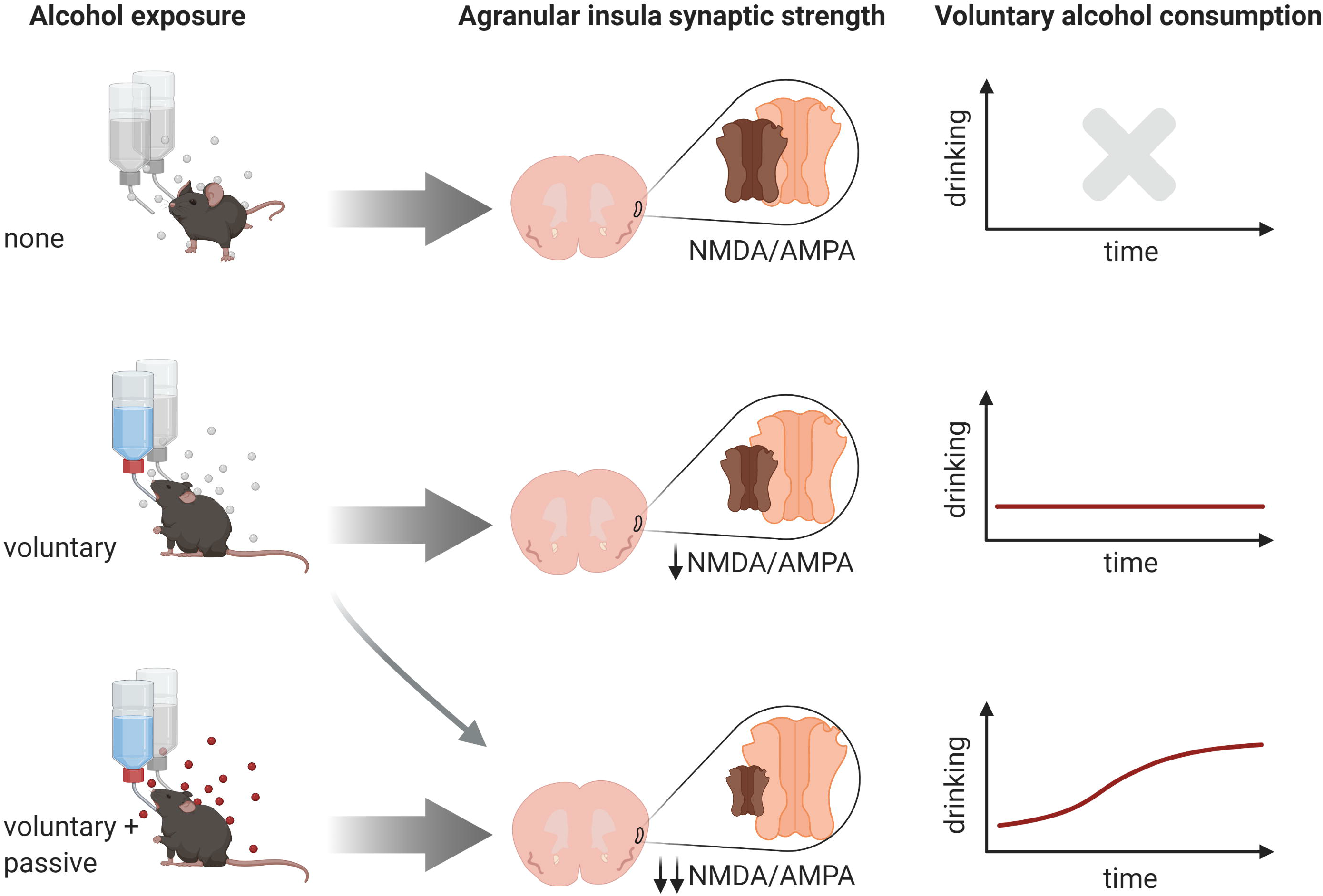
Ethanol-induced plasticity in layer 2/3 agranular insular cortex predicts escalation of ethanol consumption. Both 2BC + Air (middle) and 2BC + CIE (bottom) groups showed a suppression of the NMDA/AMPA ratio relative to ethanol-naïve controls (top). Regardless of treatment group, the NMDA/AMPA ratio in layer 2/3 pyramidal neurons was inversely related to change in drinking over time.

## Authors contribution

JS, RMa, and RMo conceived and designed experiments. JS performed the experiments. JS and RMa analyzed the data and interpreted the results. JS and RMa wrote the paper. HA and DC conducted breeding of animals and assisted with animal behavior.

## Data availability statement

The data that support the findings of this study are available from the corresponding author upon reasonable request.

## Acknowledgments

The authors wish to thank Dr. Rueben Gonzales for contributing to critical discussions of this work. The graphical abstract was created with BioRender.com. The authors declare that the research was conducted in the absence of any commercial or financial relationships that could be construed as a potential conflict of interest.

